# Multi-generational Analysis and Manipulation of Chromosomes in a Polyploid Cyanobacterium

**DOI:** 10.1101/661256

**Authors:** Kristin A. Moore, Jian Wei Tay, Jeffrey C. Cameron

## Abstract

Faithful inheritance of genetic material from one generation to the next is an essential process for all life on earth. Much of what is known about microbial DNA replication and inheritance has been learned from a small number of bacterial species that share many common traits. Whether these pathways are conserved across the great diversity of the microbiome remains unclear. To address this question, we studied chromosome dynamics in a polyploid photosynthetic bacteria using single cell, time-lapse microscopy over multi-generation lineages in conjunction with inducible CRISPR-interference and fluorescent chromosome labeling. With this method we demonstrated the long-term consequences of manipulating parameters such as cell growth, cell division, and DNA replication and segregation on chromosome regulation in a polyploid bacterial species. We find that these bacteria are surprisingly resilient to chromosome disruption resulting in continued cell growth when DNA replication is inhibited and even in the complete absence of chromosomes.

## INTRODUCTION

Strict mechanisms have been described throughout all kingdoms of life to ensure that genetic material is reliably inherited in future generations (O’Donnell, Langston, & Stillman, 2013). These mechanisms include regulating how and when DNA is replicated and the process of symmetric DNA segregation to progeny. In bacterial cells, the vast majority of the work describing DNA replication and segregation machinery has occurred in a small number of well-studied model systems (Reyes-Lamothe, Nicolas, & Sherratt, 2012). However, model systems represent only a tiny fraction of the microbial diversity, and it is now evident that this diversity plays a pivotal role in the macroscale world (Liu & Deutschbauer, 2018; Locey & Lennon, 2016). To determine whether the well-defined pathways and consequences of DNA replication and segregation are mechanistically conserved in less well studied bacterial species, we investigated chromosome dynamics in the cyanobacteria *Synechococcus* sp. PCC 7002 (hereafter PCC 7002), a polyploid photosynthetic bacteria.

Polyploidy, the presence of multiple, identical chromosome copies, is not often associated with bacteria. However, both industrially and medically relevant prokaryotes, as well as prokaryotically-derived organelles such as mitochondria and chloroplasts, are polyploid (Clay Montier, Deng, & Bai, 2009; Sakamoto & Takami, 2018; Soppa, 2017). Polyploidy is distinct from mero-oligoploidy, which occurs in rapidly growing monoploid bacteria, such as *E. coli*, when multiple rounds of DNA replication initiation begin prior to termination (Cooper & Helmstetter, 1968). This overlapping replication results in cells with an increased ratio of origin-proximal to termini-proximal DNA content (Dennis & Bremer, 2008). However, during slower growth this imbalance dissipates. In contrast, polyploid bacteria obligately contain multiple complete copies of their chromosome. DNA replication in polyploid bacterial cells appears to be stochastic and is not correlated with cell division (Chen, Afonso, Silver, & Savage, 2012; Jain, Vijayan, & O’Shea, 2012). However, the mechanisms controlling DNA copy number and segregation are not well defined.

Furthermore, the physiological consequences of polyploidy in bacteria have not been thoroughly investigated. Increasing plasmid copy number in bacterial cells results in increased gene expression (Segall-Shapiro, Sontag, & Voigt, 2018). However, the effect of increasing chromosome copy number in bacterial cells is less clear, with recent evidence indicating that growth rate and limited translational machinery are the major regulators of constitutive gene expression (Bryant, Sellars, Busby, & Lee, 2014; Chandler & Pritchard, 1975). Additional consequences of polyploidy may result from increased adaptability in extreme environments as has been shown for polyploid varieties of both plants and yeast (Selmecki et al., 2015; Van de Peer, Mizrachi, & Marchal, 2017), or as nutrient storage (Zerulla et al., 2014). Increasing our understanding of both the mechanisms and consequences of polyploidy in prokaryotes is essential to gaining a broader perspective on the microbial world and its interactions with eukaryotes and the environment.

To address these gaps in our knowledge, we fluorescently labeled chromosomes, imaged cells over multiple generations, and quantitatively analyzed parameters such as chromosome number and gene expression over time. Using this method, we define the effect of growth rate on chromosome dynamics and gene expression in PCC 7002. We also used inducible CRISPR-interference to determine the consequences of inhibiting essential cell functions, such as cell growth and division, on chromosome replication and segregation. Additionally, we demonstrate that several required regulators of DNA dynamics in monoploid cells are not essential for DNA replication and segregation in PCC 7002 and do not have consequences on cell growth. Lastly, we demonstrate that PCC 7002 is surprisingly resilitant to chromosomal insults, allowing for continued growth while DNA replication is inhibited as well as in the absence of chromosomes altogether. However, we find that irremediable chromosome loss has a dramatic effect on the photosynthetic capacity of the cell.

## RESULTS

### Chromosome labeling in PCC 7002

To study polyploidy at the single cell level in PCC 7002, we fluorescently labeled individual chromosomes using a *240x tetO* array - TetR-sfGFP (super folder GFP) based approach adapted from previously described studies in yeast and bacteria (Chen et al., 2012; Jain et al., 2012; Michaelis, Ciosk, & Nasmyth, 1997). Because this system relies on inducible TetR-*tetO* binding we were able to modulate chromosome labeling with the small molecule anhydrotetracyclene (aTC) (**Fig. S1A-B**). We also genetically encoded the *mOrange2* gene as a tool to study gene expression in these cells. To analyze chromosome number in large populations of cells we developed customized software to identify individual chromosomes, as well as measure gene expression via fluorescence intensity to generate quantitative data. **Fig. 1A** depicts representative images, from left to right, of mOrange2 expression, chromosome labeling, DNA staining (with Hoechst dye), thylakoid membrane fluorescence, and automated cell and chromosome identification masks. As previously demonstrated for other polyploid bacterial strains we observed a broad distribution of chromosome number within our population of cells (**Fig. 1B**). We also observed a strong correlation between cell length and chromosome number, as well as DNA staining intensity as measured by comparing Hoechst staining with chromosome number (**Fig. 1C-D**). Regardless of chromosome number we observe that mean mOrange2 expression (mOrange2 intensity normalized to cell area) remains consistent in all cells (**Fig. 1E**) similar to observations in a different cyanobacterial strain *Synechococcus elongatus* PCC 7942 (here after PCC 7942) (Zheng & O’Shea, 2017).

**Figure 1.**
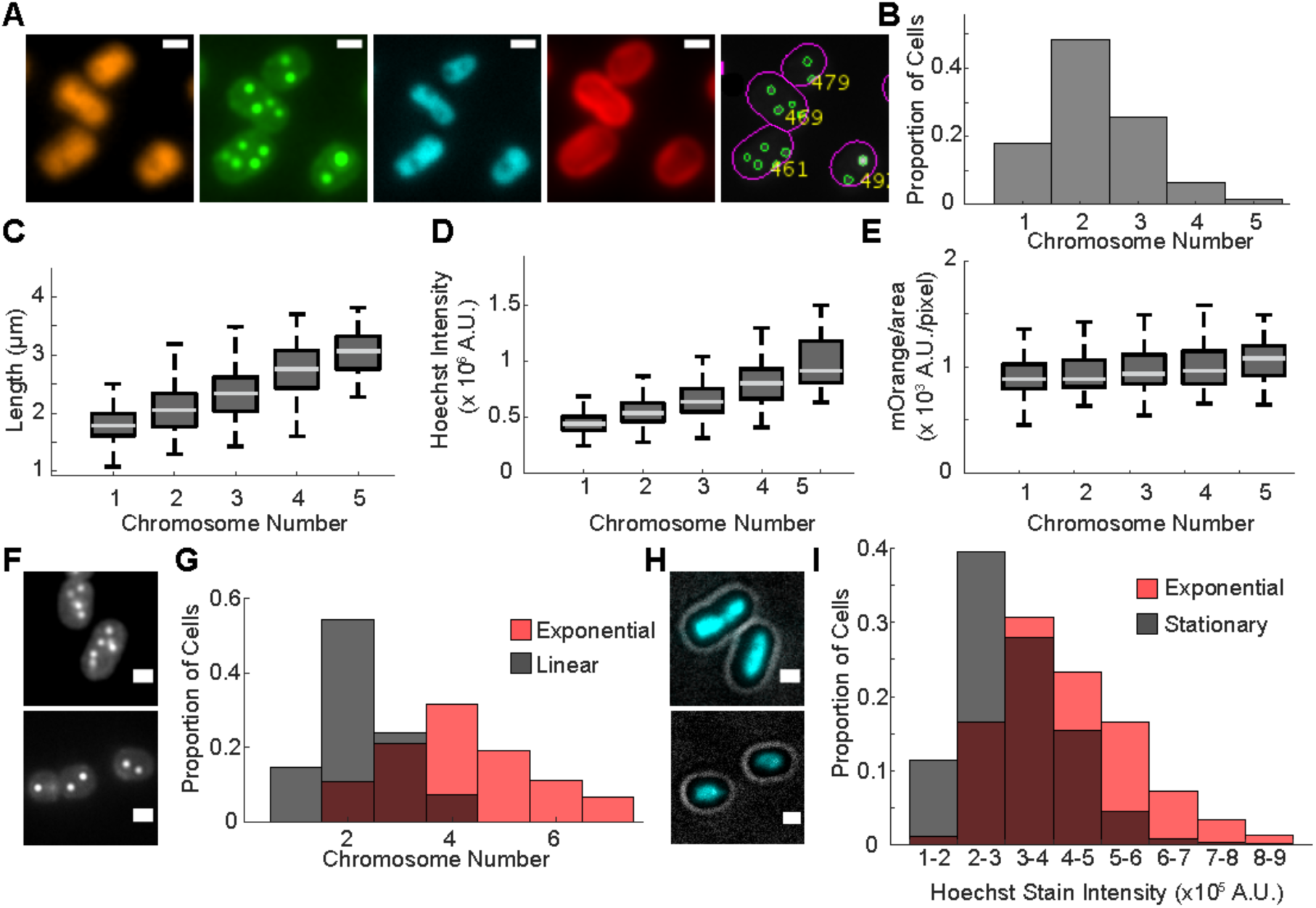
Chromosome labeling in PCC 7002. **A)** From L-R: Representative images of mOrange2 expression, sfGFP-labeled chromosomes, Hoechst staining, endogenously fluorescent thylakoid membranes, and cell (magenta outlines) and chromosome (green outlines) masks from a population of *mOrange2::240x tetO array::TetR-sfGFP* cells. **B)** The distribution of chromosome counts from the population of cells represented in Fig. 1A. **C-E)** Length, Hoechst stain intensity, and mean mOrange2 expression (mOrange2 intensity/area), respectively, measured in cells with different chromosome number from the population of cells represented Fig 1A. N = 1933. Boxes represent the 25th-75th percentile of the data, with the median marked in light gray. Whiskers show the distribution of extreme values. **F)** Representative images of chromosome labeling in cells from exponential (top panel) and linear (bottom panel) growth phase. **G)** Distributions of chromosome numbers in the populations represented in Fig. 1F. N_Exp_ = 240, N_Lin_ = 506. **H)** Representative images of Hoechst staining in cells from exponential (top panel) or stationary (bottom panel) growth phase. Hoechst stain images merged with brightfield images to denote perimeter of cell. **I)**

To determine whether growth phase has an effect on chromosome number, as observed for other polyploid bacterial strains (Soppa, 2017), we grew chromosome labeled cells to either mid-exponential or late-linear phase and imaged cells. Cells in exponential phase were both larger in size and had a higher average chromosome number than cells in linear phase (**Fig. 1 F-G**). We also grew Wild Type (WT) cells to either exponential or stationary phase and measured DNA content using Hoechst staining **(Fig. 1H)**. We observed a shift toward increased staining in exponential cells compared to stationary phase cells (**Fig. 1 H-I**), indicating that DNA content decreases in slow growing WT cells, similar to what we observe for chromosome labeled cells. We confirmed these observations using a previously described quantitative-PCR based method (**Table S1**) (Pecoraro, Zerulla, Lange, & Soppa, 2011).

Distributions of Hoechst staining divided into 8 groups by intensity values from the populations of cells represented in H. These groups are not representative of chromosome number. N_Exp_ = 967, N_Stat_ = 1089. Scale Bar = 1 µM.

### Chromosome Dynamics during PCC 7002 Growth

We captured chromosome dynamics in growing cells by performing time-lapse microscopy of chromosome labeled strains. Unless otherwise noted cells were grown on 1% (w/v) agarose pads made with A+ media under 150 µmol photons m^−2^ s^−1^ of red light (640nm) as previously described (Moore et al., 2018). Using this method, we were able to image chromosome labeled cells for ∼24 hr with a 30 minute (min) frame rate, which allowed us to track lineages for 3-4 generations (**Fig. 2A-B, Movie S1**). Using our image analysis software, we estimated both average and per frame chromosome number over time (**Fig. 2B-C**). Because of the dynamic nature of chromosome replication, which includes binding and release of DNA binding proteins such as TetR, as well as limitations of our frame rate, the number of puncta in a cell may not represent the true number of chromosomes in that cell. To address this challenge, we created an algorithm to predict chromosome number based on previous and future frames for each cell. (**Fig. S2** and methods section for details). These corrected values were used for all data analysis. Unless otherwise noted, only cells with traces that both started and ended during the imaging time (i.e. not first or last generation cells) were analyzed.

**Figure 2.**
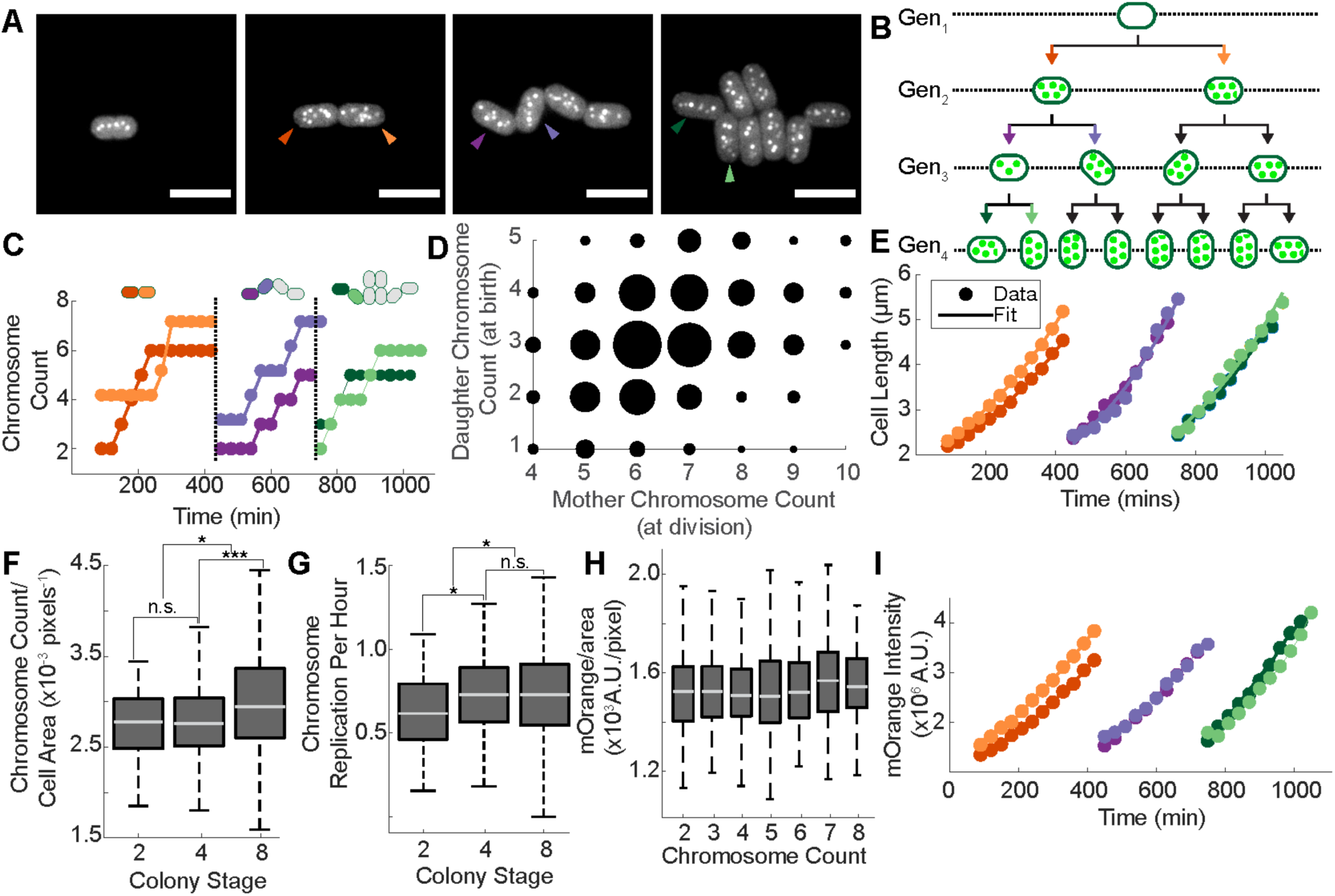
Chromosome dynamics in PCC 7002. **A)** Time-lapse images of labeled chromosomes from *mOrange2::tetO array:: TetR-sfGFP* cells grown in 150 µmol photons m^−2^ s^−1^ red light for 18.5 hours (hr). Colored arrows correspond to the colored traces in throughout Fig. 2. **B)** Average chromosome number calculated for cells at each microcolony position and stage. **C)** Chromosome count over time for cells at the 2-cell (orange), 4-cell (purple) and 8-cell (green) colony stages. Only two representative traces are displayed for the 4- and 8-cell stages for the sake of clarity. Dotted lines indicate cell division events. **D)** Chromosome segregation from mother cells (chromosome count at division on X-axis) to daughter cells (chromosome count at birth on Y-axis). Spot size is proportional to the number of cells. N_mother_ = 385, N_daughter_ = 770. **E)** Cell length over time for cells at different growth stages of microcolony formation. An exponential model was used for length doubling time calculations, displayed as solid lines. **F-G)** Chromosome count to cell area ratio, averaged over cell lifespan, and chromosome replication rate, respectively, for all cells in the 2-, 4-, and 8-cell microcolony stages. N_2-cell_ = 64, N_4-well_ = 229, N_8-cell_ = 332. **H)** mean mOrange2 intensity for cells with different chromosome numbers at the 4-cell microcolony stage. N = 2488 single frames. **I)** Total mOrange2 intensity measured over time for cells at the 2-, 4-, and 8-cell stages. Unless otherwise noted scale bar = 5 µm in all figures.

Using this platform, we were able to analyze chromosome dynamics in a polyploid bacterial strain over multi-generational lineages for the first time. One of the most striking observations from our movies was the lack of regularity in both chromosome number and replication. Although we do not have the temporal resolution to determine precise chromosome numbers at all times, we are able to visualize distinctly different patterns of chromosome replication throughout our time-lapse images. Replication appears to be continuous throughout the cell cycle and does not pause before cell division (**Fig 2C, Movie S1**). The average chromosome number over a cell lifespan varied between 2.6-7.2 with a median value of 4.7, and did not appear to depend on microcolony position (**Fig. 2B**). Consistent with previously observed data (Chen et al., 2012; Jain et al., 2012), we note that chromosomes segregate relatively evenly during division, but that there is some variation. Furthermore, the absolute number of chromosomes at division is not constant, and not always an even number, indicating that daughter cells within a population, even those from the same mother cell, may not inherit the same amount of genetic material **(Fig 2D)**. We did not observe specific positioning of chromosomes prior to division as was observed for in PCC 7942, but did see relatively even spacing of chromosomes throughout the cell (Chen et al., 2012).

We calculated metrics such as growth rate for each cell (**Fig. 2E**). Cells with labeled chromosomes grew similarly to WT cells as noted by the characteristic microcolony formation of PCC 7002 (Moore et al., 2018). Measured growth rates were also similar in chromosome labeled and WT cells, with median length doubling times of 266 and 245 min, respectively. Chromosome number to cell area ratio remained relatively constant for cells in different stages of microcolony formation, with a slight, but statistically significant, increase in the ratio at the 8-cell stage compared to the 2- and 4-cell stages (**Fig. 2F**). We also observed that the rate of chromosome replication, measured by dividing the difference between starting and ending chromosome number in each cell by the amount of time between cell birth and division, was similar in different stages of microcolony growth with a small decrease in the 2-cell stage compared to the 4- and 8-cell stages (**Fig. 2G**). As we and others have observed in non-time lapse imaged cells (Zheng & O’Shea, 2017), mean protein expression was similar during growth for cells with different numbers of chromosomes under these conditions (**Fig. 2H**). Interestingly, we did not detect differences in either the total amount or rate of mOrange2 accumulation for cells with differential chromosome replication patterns (**Fig. 2I**), indicating that variability in chromosome copy number does not lead to variability in expression of constitutively expressed proteins under these conditions.

### Growth Rate Correlates with Chromosome Replication Rate and Gene Expression Changes

Because PCC 7002 can use photosynthesis as its sole energy source, we were able control growth rate by varying the growth light intensity. To determine how changing growth conditions affects chromosome replication and inheritance, we grew cells with 45 µmol photons m^−2^ s^−1^ of red light rather than the 150 µmol photons m^−2^ s^−1^ described above. Cells still formed similar microcolony formations during growth (**Fig. 3A**), but on a longer time scale (**Fig 3B and F**). We again observed variability in the patterns of chromosome replication (**Fig. 3C**). However, the distributions of chromosome to area ratio calculated for cells grown in high light (150 µmol photons m^−2^ s^−1^ - HL) or low light (45 µmol photons m^−2^ s^−1^ - LL) are indistinguishable (**Fig. 3D**) indicating that chromosome replication must proportionally decrease in slower growing cells. This is confirmed by the significantly slower chromosome replication rate, median value of 0.33 chromosomes replicated per hour, measured in cells growing in LL conditions.

**Figure 3.**
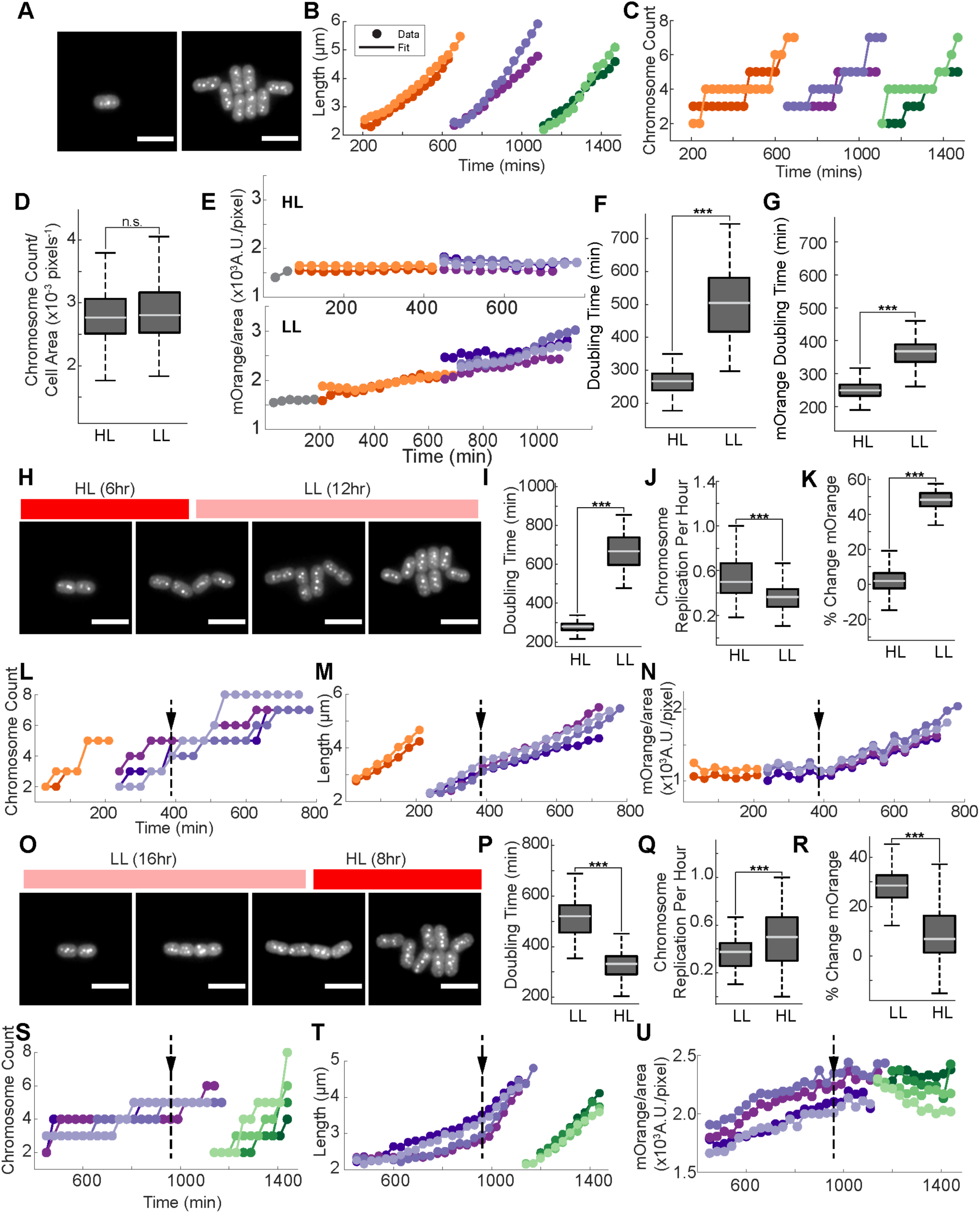
Effect of growth rate on chromosome dynamics and gene expression. **A)** Time-lapse images of labeled chromosomes from *mOrange2::tetO array::TetR-sfGFP* cells grown in 45 µmol photons m^−2^ s^−1^ red light over 27 hr. **B-C)** Length and chromosome count, respectively, at different stages of microcolony growth. **D)** Chromosome count per area ratio, averaged over the lifespan of cells, grown in HL or LL. **E)** mean mOrange2 intensity over time for cells grown in HL (top panel) or LL (bottom panel). **F-G)** Cell length doubling time and mOrange2 intensity doubling time, respectively, for cells grown in HL and LL. Data from 4-cell stage displayed. N_HL_ = 229, N_LL_ = 216 for Fig. 3D, F, and G. **H)** Time-lapse images of labeled chromosomes in cells grown in HL for 6 hr followed by LL for 12 hr. **I-K)** Length doubling time, chromosome replication rate, and percentage change in mOrange2 intensity, respectively, for cells grown in HL and LL. Percentage change was measured by dividing the difference between mOrange2 intensity in the final and the first frame of a cell trace by the initial intensity, multiplied by 100. N_HL_ = 98, N_LL_ = 79. **L-N**) Chromosome count, length, and mean mOrange2 intensity, respectively, over time for cells grown in HL (2-cell stage, orange traces) or during the light transition (4-cell stage, purple traces), indicated by the arrow and dotted line. **O)** Time-lapse images of labeled chromosomes in cells grown in LL for 16 hr followed by HL for 8 hr. **P-R)** Length doubling time, chromosome replication rate, and percentage change in mOrange2 intensity, respectively, for cells grown in LL and HL. N_LL_ = 77, N_HL_ = 77. **S-U)** Chromosome count, length, and mean mOrange2 intensity, respectively, over time for cells grown in during the light transition (4-cell stage, purple traces) and in HL (8-cell stage, green traces, only 4 traces shown for clarity).

Interestingly, mean mOrange2 intensity increased in cells grown at low light (**Fig. 3E-Bottom Panel**), resulting in a two-fold increase in mOrange2 intensity at the end of four-cell stage of growth compared to starting mOrange2 intensity. In comparison, mOrange2 intensity for cells grown in HL did not increase over the same cell-stages (**Fig. 3E-Top Panel**). To determine whether the rate of mOrange2 gene expression increased in LL conditions, we calculated the amount of time it took for cells to double their mOrange2 intensity when grown in either HL or LL (**Fig. 3G**). Slower growing cells have longer mOrange2 doubling times compared to fast growing cells indicating a *decrease* in gene expression in LL conditions. However, the differences in gene expression rate were smaller than the differences in physical growth rate; in LL, cells expressed less mOrange2 but accumulated more of it over their lifespan (compare doubling times **Fig. 3F and 3G**).

Decreases in chromosome replication rate and growth rate appear to occur almost instantaneously when cells are shifted from HL to LL, as changes are evident in the first hour after light conditions have decreased (**Fig 3L-M**-dashed line notes first frame after light shift). Interestingly, mOrange2 intensity also starts to increase as soon as cells are shifted to LL, further supporting the hypothesis that changes in growth rate rather than gene expression (which is expected to have a delayed effect) are responsible for increased accumulation of mOrange2 in these cells (**Fig. 3N**). After shifting cells from LL to HL, changes in chromosome replication are apparent in cells grown entirely in HL, but these changes are not observed in cells that span the light transition (**Fig. 3S, purple traces**). Increases in growth rate and decreases in mOrange2 intensity are observed immediately after the LL to HL transition (**Fig. 3T-U**) indicating that these responses may occur on a faster time scale than regulation of chromosome replication for this light transition.

### Chromosome Replication is not Dependent Cell Division or Extension

Because many of the genes known to affect cellular processes such as DNA replication and segregation are essential for survival, we modified the CRISPR interference (CRISPRi) system described for PCC 7002 by Gordon et al., (2016) to make an IPTG inducible guide RNA (sgRNA), and engineered it into a strain expressing dCas9, TetR-sfGFP, and the *tetO* array. This system allowed us to temporally control essential gene expression. We grew cells in an uninduced state followed by addition of IPTG immediately prior to time-lapse imaging. With these strains we were able to observe the dynamics of chromosome replication and division as essential gene expression decreased over time.

Previous studies have noted that inhibiting cell division in monoploid bacteria does not immediately impact nucleoid number or spacing (Dai & Lutkenhaus, 1991). However, upon prolonged blockage of cell division, DNA replication is inhibited (Arjes et al., 2014). To determine whether cell division regulates DNA replication in a polypoid bacterial species, we imaged cells for 20hr after induction of an sgRNA targeting *ftsZ*, an essential gene for cell division (**Fig. 4A, Movie S2**). Uninduced cells grew comparably to cells lacking a guide RNA. However, within one generation of *ftsZ* CRISPRi induction, we observed extensive elongation of cells as well as continuous chromosome replication (**Fig. 4B-C**). The distribution of chromosome number to cell area ratios was marginally higher in *ftsZ* CRISPRi induced cells indicating the blocking cell division does not have a large effect on chromosome dynamics (**Fig. 4D**). Over the 20 hr imaging span (approximately the equivalent of five uninduced doublings), we did not observe any decrease in cell growth (**Fig. 4B**). Unfortunately, due to focusing issues of large cells growing on top of each other, it became impossible for us to consistently identify chromosomes in our images beyond the 14 hr mark of our movie for most of our cell traces. However, it does not appear that chromosome replication is inhibited by FtsZ depletion (**Fig. 4E, Movie S2**), nor do we see any irregular patterns of DNA localization as previously observed in *Bacillus subtilis* and *Staphylococcus aureus* (Arjes et al., 2014). These results indicate that unlike several model monoploid bacterial strains, chromosome replication and cell division are essentially uncoupled in polyploid bacteria.

**Figure 4.**
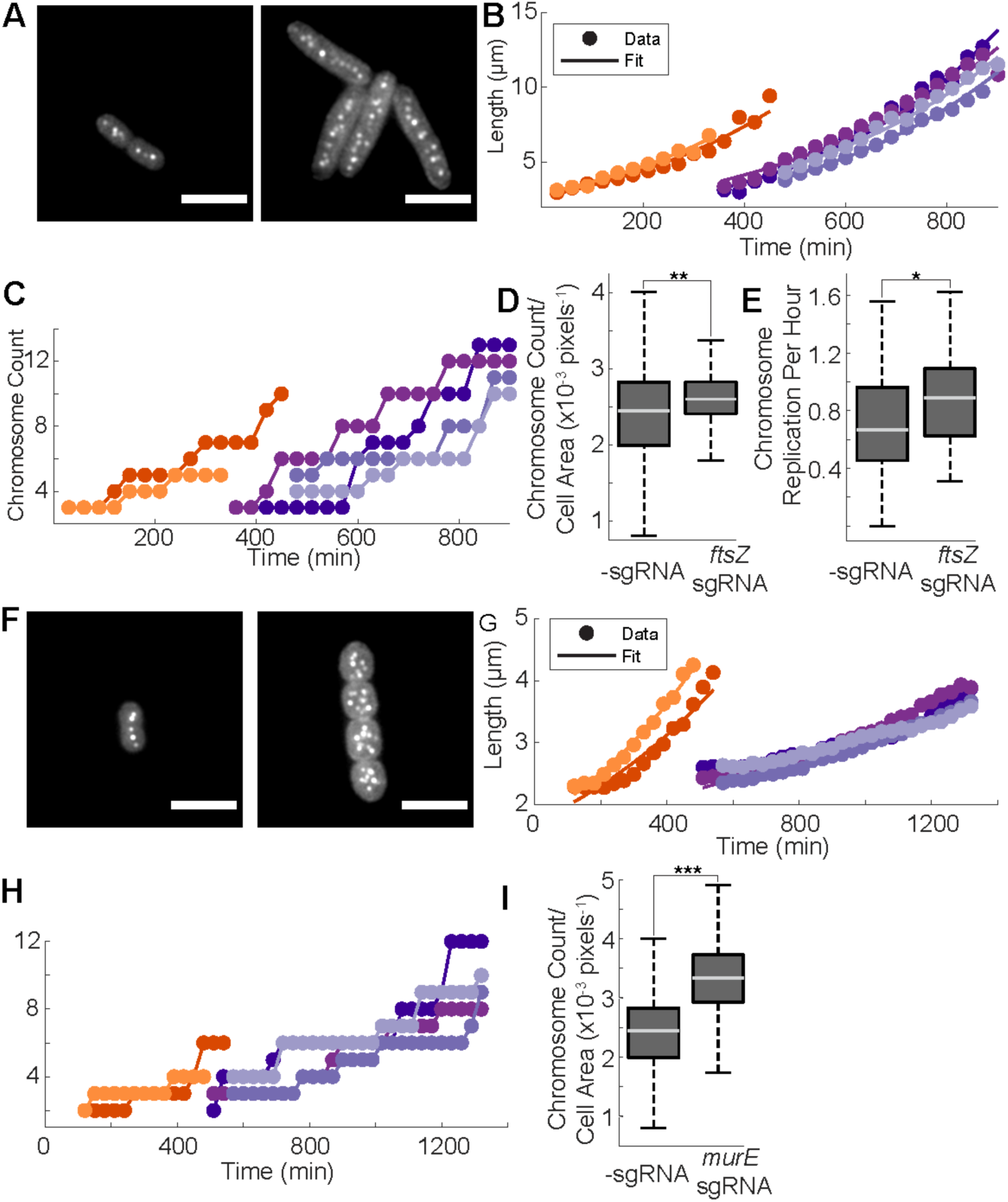
The effect of cell division and elongation rate on chromosome dynamics. **A)** Time-lapse images of labeled chromosomes in *ftsZ sgRNA::tetO array::TetR-sfGFP-dCas9* (CRISPRi) cells grown for 21.5 hr. For all CRISPRi time-lapse images, 5mM IPTG was added to agarose imaging pad to induce sgRNA expression 15 minutes prior to the first image. **B-C)** Cell length and chromosome count, respectively, in 1st (orange) and 2^nd^ (purple) generation *ftsZ*-targeting CRISPRi cells. **D-E)** Chromosome count per cell area averaged over the lifespan of the cell and chromosome replication rate, respectively, for cells with or without an sgRNA targeting *ftsZ* (2^nd^ generation and later). N_-sgRNA_ = 315, N_ftsZ CRISPRi_ = 93. **F)** Time-lapse images of labeled chromosome from *murE sgRNA::tetO array::TetR-sfGFP-dCas9* (CRISPRi) cells grown for 21.5hr. **G-H)** Cell length and chromosome count, respectively, in 2^nd^ (orange) and 3^rd^ (purple) generation *murE*-targeting CRISPRi cells. **I)** Chromosome count per cell area averaged over the lifespan of the cell for cells with or without a sgRNA targeting *murE* (third generation and later). N_-sgRNA_ = 315, N_murE CRISPRi_ = 307.

To test how cell growth and cell shape affect chromosome replication and segregation, we knocked down *murE*, an essential peptidoglycan synthesis gene. Loss of MurE from *B. subtilis* results in short round cells that are unable to form typical oblong (colloid) shapes (Peters et al., 2016). After induction of *murE* CRISPRi, we observed similar morphological defects to those previously described, as well as cells that elongate at an extremely slow rate (**Fig. 4F-G, Movie S3**). When growth rate was genetically manipulated, we observed an accumulation of chromosomes in fully induced cells, as well as a significantly increased chromosome to cell area ratio for later generation cells (**Fig 4H-I**). These results indicate that cell elongation is not essential for chromosome replication.

### Inhibiting chromosome Replication Stalls Cell division, but not Growth

Previous work in several polyploid prokaryotes has indicated that current models of DNA replication, specifically the requirement for *dnaA* and a well-defined *oriC* site, may not be conserved across the microbiome (Gehring et al., 2017; Ohbayashi et al., 2016; Richter, Hagemann, & Messer, 1998). In support of this assertion, we found that in PCC 7002, *dnaA* (*Synpcc7002_A0001*) was not required for DNA replication for any conditions tested (**Fig. 5A-B**). *ΔdnaA* cells with labeled chromosomes grow and have similar chromosome to area ratios as cells that are WT at the *dnaA* locus under standard growth conditions (**Fig 5C-D**). *dnaA* deletion was confirmed by PCR (**Fig. S3A**). There are no additional *dnaA* homologs within PCC 7002, and canonical clustering of DnaA binding sites is also absent from the genome, further indicating that the role of DnaA is not conserved across prokaryotic species.

**Figure 5.**
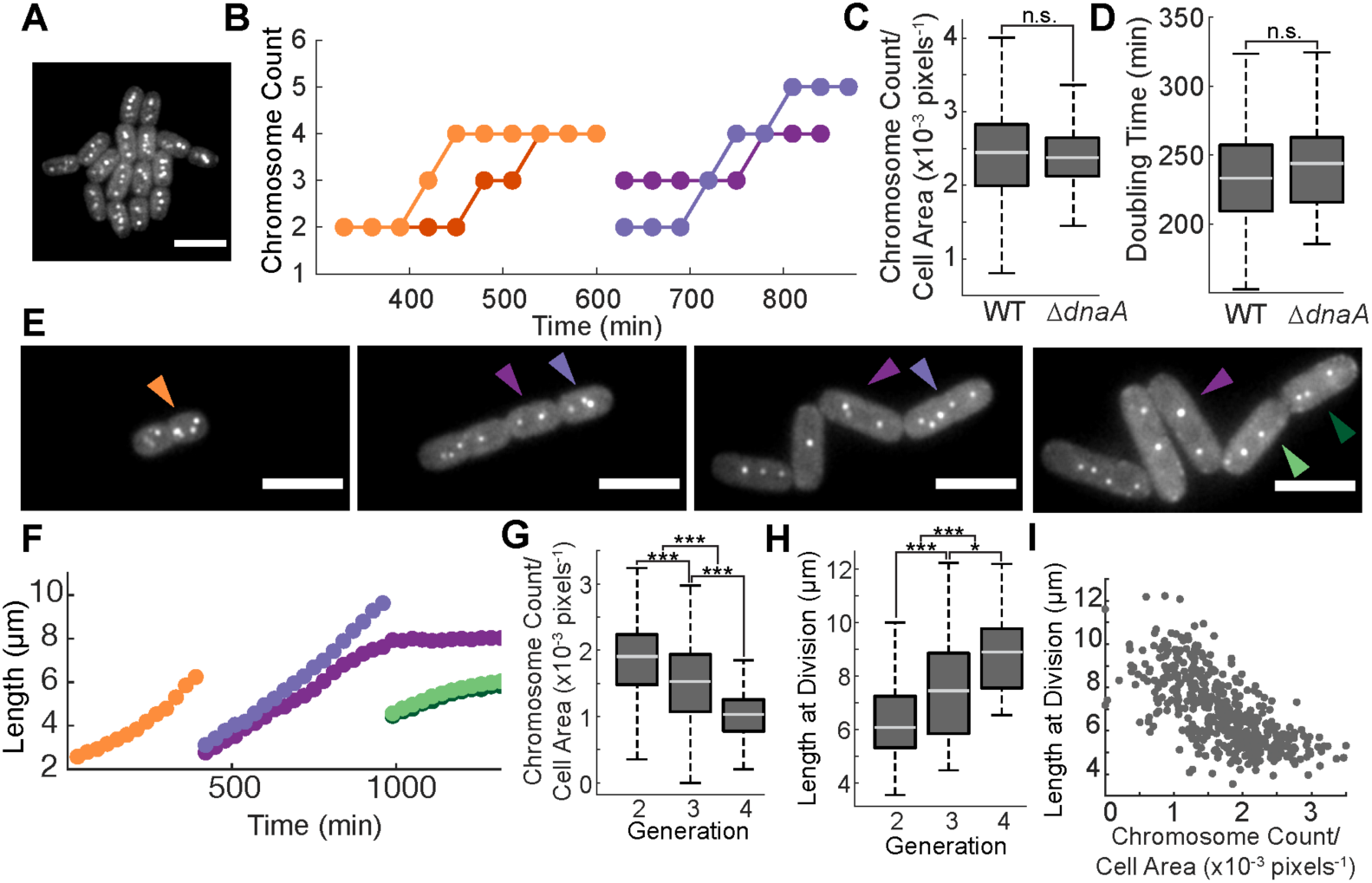
Chromosome replication in PCC 7002. **A)** Labelled chromosomes at the 16-cell microcolony stage in *ΔdnaA::TetO array::TetR-sfGFP* cells. Cells grown for 19.5 hr. **B)** Chromosome count over time at the 2-cell (orange) and 4-cell (purple) stage of *ΔdnaA::TetO array::TetR-sfGFP* cells. **C-D)** Chromosome count per cell area averaged over the lifespan of the cell and length doubling time, respectively, for cells WT or mutant at the *dnaA* locus. N_WT_ = 272, N*_ΔdnaA_* = 124 **E)** Time-lapse images of *dnaX sgRNA::tetO array::TetR-sfGFP-dCas9* CRISPRi cells grown for 26 hr. Arrow colors correspond to graphs in F. **F)** Length over time for 1st (orange), 2^nd^ (purple), and 3^rd^ (green) generation *dnaX* CRISPRi cells. **G-H)** Chromosome count per cell area averaged over the lifespan of the cell, and cell length at division for 2^nd^, 3^rd^, and 4^th^ generation *dnaX* CRISPRi cells, respectively. **I)** Scatter plot of length at division and chromosome count per cell area in *dnaX* CRISPRi treated cells. N = 470.

To inhibit DNA replication, we used CRISPRi to target *dnaX*, an essential component of the DNA polymerase holoenzyme (Blinkova et al., 1993). This method allowed us to visualize dilution of chromosomes over time (**Fig 5E, Movie S4**). Initial inhibition of DNA replication did not affect cell growth (**Fig. 5F**-compare initial length measurements between 1^st^ and 2^nd^ generation cells) even though chromosome to area ratio had already dropped significantly lower than -sgRNA cells by the 2^nd^ generation (compare **Fig. 5G** and **Fig. 2F**). As chromosome depletion continued, cell division was inhibited, resulting in cells that either did not divide or were delayed in division, leading to significantly longer cells. These trends became more pronounced in later generations (**Fig. 5G-H**). We observed an inverse correlation between cell length at division and chromosome per area ratio (**Fig. 5I**), indicating that DNA content per cell area plays a pivotal role in cellular homeostasis. Whether these effects are driven by changes to gene expression and protein concentration or other undefined checkpoints on cell division, remains an open question in PCC 7002. Interestingly, chromosome localization in these cells appears regulated as even minimal numbers of chromosomes remain relatively evenly spaced throughout each cell (**Fig. 5E**-right-most panel).

### DNA Segregation in PCC 7002 is Not Driven by Known Mechanisms

Because DNA segregation into daughter cells is essential for cell viability, bacteria have evolved mechanisms to ensure faithful segregation of both chromosomes and plasmids in the next generation (Wang, Llopis, & Rudner, 2013). To determine whether the well-defined ParABS system, which is essential for chromosome and plasmid segregation in many bacterial strains, affects chromosome segregation in PCC 7002, we created a *ΔparA* strain by deleting the chromosomal copy of *parA (Synpcc7002_A0432). parA* deletion was confirmed by PCR (**Fig. S3B**). *ΔparA* cells did not appear to have any growth defects, nor chromosome segregation errors (**Fig 6A-B**). Additionally, we were not able to visualize the chromosomal homolog for ParB (*Synpcc_A0433*) after tagging with sfGFP, which has been used as a method for chromosome labeling in other bacteria (Lee, Lin, Moriya, & Grossman, 2003). These results are similar to observations made in *Synechococcus elongatus* PCC 7942 (Jain et al., 2012), indicating that the ParABS system does not play a role in chromosome segregation in polyploid cyanobacteria.

**Figure 6.**
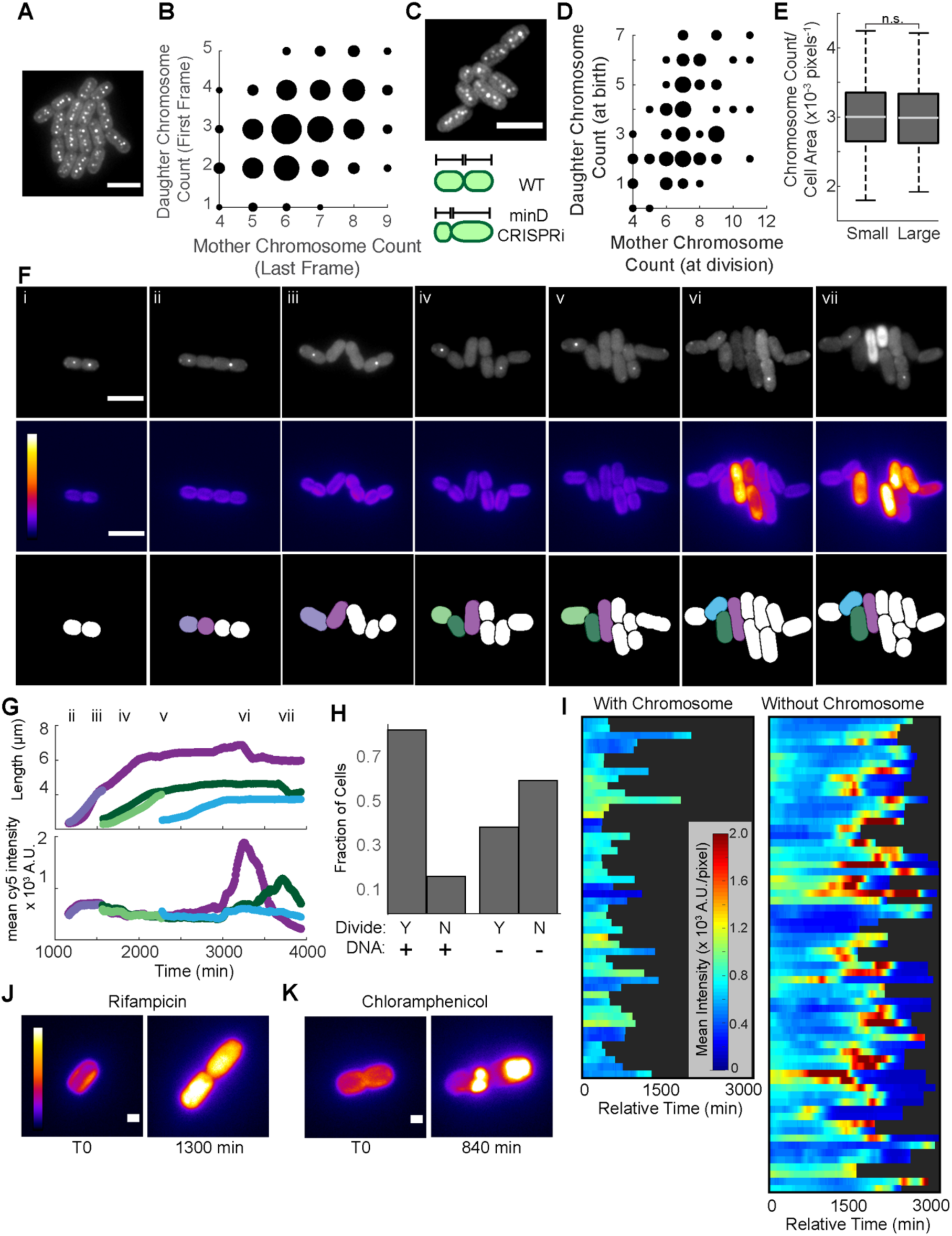
Chromosome Segregation in PCC 7002. **A)** Labelled chromosomes at the 16-cell microcolony stage in *ΔparA::TetO array::TetR-sfGFP* cells. Cells grown for 20 hr. **B)** Chromosome segregation of *ΔparA::TetO array::TetR-sfGFP*. Nmother = 502, Ndaughter = 1004. **C)** Top panel: 8-cell microcolony of *minD sgRNA::TetO array::TetR-sfGFP-dCas9*. Bottom panel: Schematic of the effect of MinD depletion on cell division. **D)** Chromosome segregation plot of *minD sgRNA::TetO array::TetR-sfGFP-dCas9*. Only cells in the 3^rd^ generation or later were analyzed. N_mother_ = 43, N_daughter_ = 86. **E)** Chromosome Count to cell area ratio distributions for small cells (starting cell length < 10th percentile of -sgRNA cells) and large cells (starting cell length > 90th percentile of -sgRNA cells). N_S_ = 125, N_L_ = 175. **F)** Time-lapse images of labeled chromosomes (top panel) and endogenous fluorescence from thylakoid membranes (middle panel) without aTC for 65 hr. Bottom panel represents cell masks with colors corresponding to cell traces in G. Calibration bar represents intensities between 75-2450 A.U. **G)** Cell length (top panel), and mean emission intensity after 640 nm excitation representing photosynthetic capacity (bottom panel), plotted over time. **H)** The fraction of cells with at least one (DNA+) or without chromosomes (DNA-) that divide within 25hr. N = 310. **I)** Heatmap of mean emission intensity after 640nm excitation over time for cells with (left) or without (right) a chromosome spot. T0 = first frame after division. Cells with only one chromosome spot that divided and cells without chromosomes that did not divide were analyzed. N = 116. **J)** Time-lapse images of *mOrange2::tetO array:: TetR-sfGFP* cells treated with rifampicin to block transcription. **K)** Time-lapse images of *mOrange2::tetO array::TetR-sfGFP* cells treated with chloramphenicol to block translation. For J and K, scale bar represents 1µm.

We also created a CRISPRi line targeting *mreB*, another known regulator of chromosome segregation (Gitai, Dye, Reisenauer, Wachi, & Shapiro, 2005; Kruse et al., 2006). *mreB* does appear to be essential in our cells, but not due to defects in chromosome segregation. MreB depleted cells grew into 3-dimensional spheres that were very fragile and often burst during imaging, indicating that cell structure defects associated with loss of MreB leads to defects in viability (**Movie S5**). Although we were unable to count chromosomes in these cells due to their 3-dimensional shape, we did not observe any obvious defects in chromosome segregation, indicating that MreB does not play a role in chromosome segregation in PCC 7002. Our results are consistent with other studies of that have demonstrated a structural function for MreB in polyploid cyanobacteria, but not a role in chromosome segregation (Hu, Yang, Zhao, Zhang, & Zhao, 2007).

To determine whether an undefined mechanism of strict chromosome segregation exists in PCC 7002, we depleted *minD* with CRISPRi. Cells depleted of MinD have irregular septum placement, resulting in both larger and smaller cells than average in -sgRNA strains (**Fig. 6C**). As evident from both visual and quantitative analysis, it is clear that chromosomes were not split evenly into large and small cells, but rather that the number of chromosomes segregated into each daughter cell was proportional to cell size (**Fig. 6D, Movie S6**). MinD depleted cells provided us with an additional tool to study differences between large and small cells with very different absolute chromosome numbers. When we grouped cells by cell length at birth, we did not observe distinct differences in cell physiology (**Movie S6**). The number of chromosomes per area were also almost identical for the smallest and largest cells (**Fig. 6E**). These results indicate that chromosome segregation in PCC 7002 may occur randomly based on chromosome localization and septum placement.

### DNA Segregation Errors Affect Photosynthetic Activity

Previous work with the TetR-*tetO* array system has indicated that strong binding of TetR to the array can lead to chromosome replication and segregation errors in monoploid bacteria (Bernard, Marquis, & Rudner, 2010). Under standard growth and imaging procedures, we grew cells with aTC to ensure that chromosome dynamics were minimally perturbed. However, to determine how excessive TetR binding affected PCC 7002, we grew chromosome labeled cells to stationary phase without aTC in shaking flasks and imaged cells without aTC over a 65 hr time frame. Under these conditions, we observed cells with single GFP-labeled puncta, which may represent one or more chromosomes (**Fig. 6F**). Unlike stationary phase cells grown in the presence of aTC (**Movie S7**), chromosome number did not increase with growth in the vast majority of cells when grown without aTC (**Movie S8**). This may be representative of a block in chromosome replication, oligomerization of sfGFP bound to multiple chromosomes, or both.

Interestingly, cells with single chromosome spots readily grew and divided, resulting in uneven segregation of the chromosomes between daughter cells (**Fig. 6F, Movie S8**). Surprisingly, cells that did not receive a chromosome after division continued to grow (**Fig. 6G**, dark purple and dark green traces), and ∼40% of these cells were able to divide at least once, indicating that the absence of a chromosome has no effect on initial cell growth, and that neither the presence of a chromosome nor its replication is absolutely required for cell division in PCC 7002 (**Fig. 6H**). After ∼1-2 cell length doubling periods, growth arrest occurs in cells lacking chromosomes. After ∼24 hr, we observed an increase in endogenous fluorescence originating from the thylakoid membranes, evidenced by the increased emission after 640 nm excitation, indicating that the photosynthetic machinery had become dysfunctional (Maxwell & Johnson, 2000) (**Fig 6F** middle panel, **G** bottom panel **and I** right panel). Endogenous fluorescence has been pseudocolored in **Fig 6F** to better visualize changes in intensity. In contrast, endogenous fluorescence did not increase in cells with at least one chromosome spot (**Fig 6I**-Left panel). The consistency of the timing between chromosome loss and spikes in endogenous fluorescence was relatively regular, indicating that cells have the capacity to grow, divide, and perform photosynthesis for a surprisingly long period of time without genetic inputs (**Fig 6G**).

To determine whether inhibiting gene expression through transcriptional or translational inhibition resulted in similar effects on photosynthetic machinery, we treated cells with either rifampicin to block transcription, or chloramphenicol to block translation. We observed similar increases in endogenous fluorescence with both inhibitors after slightly different time periods, which may represent the temporal effects of blocking different steps in the gene expression pathway (**Fig. 6J-K**).

## DISCUSSION

Although the regulation of DNA replication and inheritance have been long studied in model strains of monoploid bacteria, our results raise questions about our fundamental understanding of these processes across the microbiome. Here, we used single-cell time-lapse imaging to demonstrate both conserved and variable characteristics of bacterial DNA regulation between traditionally studied monoploid bacteria and polyploid strains. Our data support the notion that the ratio of chromosome content to cell size is strictly preserved under changing growth conditions as observed in monoploid strains (Donachie & Begg, 1989; Sargent, 1975). However, chromosome replication does not depend on, nor does it cause cell growth or division, as evidenced by cells accumulating chromosomes when growth is inhibited and diluting out chromosomes when DNA replication is initially stalled. Significant inhibition of DNA replication or complete chromosomal loss eventually inhibits both cell growth and division, indicating that downstream effects of chromosome replication do affect cell viability. These results support a model in which DNA replication and segregation are independent of cell growth and division in polyploid cells.

How chromosome replication and segregation are regulated in PCC 7002 remains an open question. DnaA, which is required for DNA replication in all monoploid strains studied to date, does not play an essential role in DNA replication in PCC 7002. DnaA has been shown to be dispensable for DNA replication in some, but not all, polyploid cyanobacteria and archaea stains (Gehring et al., 2017; Ohbayashi et al., 2016), indicating that polyploidy alone does not negate the need for DnaA in DNA replication. Additionally, while chromosome localization within PCC 7002 appears to be important for the proper segregation of chromosomes into daughter cells, the mechanisms driving chromosome localization are not clear. Previously described proteins involved in chromosome segregation, such as the ParABS system and MreB, are not required in PCC 7002. Alternative hypotheses based on physical parameters that mediate chromosome positioning (Jun & Wright, 2010) may play a role in chromosome localization in WT cells but do not explain why chromosomes remain evenly spaced when chromosome number has been depleted (**Movie S4**). Our data indicate that chromosome positioning is a regulated process, but the mechanisms mediating this regulation remain unknown.

In a move towards determining the consequences of polyploidy on cell physiology, we monitored constitutive gene expression in polyploid cells. We find that chromosome copy number is not a major factor in determining protein content in polyploid bacteria. Rather, growth rate appears to be the predominant driver of protein accumulation in these cells. We observe that shifts in growth rate and constitutive gene expression occur very rapidly within the same generation of environmental changes, indicating that these responses likely do not depend on transcriptional or translational “re-programming” of the cell. Gene expression in these experiments was driven by a non-native promoter, which may respond differently than endogenous systems. However, our results support the notion that DNA content is often not the limiting factor in protein production, and are consistent with predictions made by several recent models of gene expression (Klumpp, Zhang, & Hwa, 2009; Lin & Amir, 2018). In polyploid cells, it is possible that heterozygous alleles may be present on different chromosomes. It will be interesting for future studies to determine whether copy number may in fact be important for relative gene expression from allelic variants in polyploid cells. A more thorough understanding of how chromosome copy number and growth rate affect nuanced types of gene expression, such as inducible and native promoter systems, in both monoploid and polyploid cells, will be essential for understanding general bacterial physiology, as well as for designing functional synthetic biology circuits in bacteria.

Lastly, we demonstrated that polyploid cells are surprisingly robust to chromosomal insults and loss. Cells lacking chromosomes are able to grow, and almost half are even able to divide, indicating that cell cycle specific gene expression is not required for division in PCC 7002. Eventually, photosynthetic damage is observed, but not for multiple cell doubling periods. We predict that photosynthetic damage results from the loss of gene expression, as similar effects on photosynthetic capacity were observed when transcription and translation were inhibited. Our data are consistent with previous studies demonstrating that gene expression is known to be essential for photosystem II repair after either visible or UV light mediated damage, and that substantial DNA damage also results in deficient photosystem damage repair (Aro, Virgin, & Andersson, 1993; Sass, Spetea, Máté, Nagy, & Vass, 1997; Vass, Kós, Sass, Nagy, & Vass, 2013). These observations indicate that polyploid organisms may be especially resistant to chromosomal insults allowing them time to attempt remediation before loss of viability. Because of this resilience, it is possible that these “chromosome-less” cells could function as photosynthetically active proto-cells and be useful tools for creating synthetic organisms in the future.

## ACKNOWLEDGEMENTS

We thank the entire Cameron Lab for thoughtful insights and critical evaluation of this work as it was being prepared. We specifically thank Patrick Thomas for reviewing and editing this manuscript. We thank the Laboratory for Interdisciplinary Statistical Analysis (LISA) at CU Boulder for advice on the appropriate statistical tests to use with our data. We also thank Brian Pfleger’s lab for providing PCC 7002 cells. Funding: This research was supported by the NSF Postdoctoral Research Fellowship in Biology under Grant No.1711932 (to K.A.M.). All opinions expressed in this paper are the authors’ and do not necessarily reflect the policies and views of NSF.

## AUTHOR CONTRIBUTIONS

Conceptualization, J.C.C.; Methodology, J.C.C. and K.A.M; Software, J.W.T; Validation J.W.T. and K.A.M; Formal analysis, Investigation, and Writing – Original Draft, K.A.M; Writing – Review and Editing, J.C.C, J.W.T, and K.A.M; Visualization, J.C.C., J.W.T., and K.A.M., Funding, J.C.C. and K.A.M; Supervision, J.C.C and K.A.M.

## METHODS

### Contact for Reagent and Resource Sharing

Further information and requests for resources and reagents should be directed to, and will be fulfilled by, the Lead Contact, Jeffrey Cameron.

### Experimental Model and Subject Details

All Strains of *Synechococcus* sp. PCC 7002 were cultivated in A+ media (Stevens, Patterson, & Myers, 1973b) in an AL-41L4 Environmental Chamber (Percival Scientific, Perry, IA) maintained at 37 ° C with atmospheric CO_2_ conditions, with continuous illumination (∼150 µmol photons m^−2^ s^−1^) provided by cool white fluorescent lamps. All strains were grown in 25 ml liquid cultures in baffled flasks (125 ml) contained with a foam stopper (Jaece identi-plug) and on an orbital shaker (200 rpm), or on medium solidified with Bacto Agar (1%; w/v). For maintenance, all *240x tetO-array:TetR-sfGFP* strains were grown with 0.5µg/mL anhydrotetracycline (aTC). Antibiotics were provided to solid medium for routine maintenance of mutants when necessary (km, 30 µg/ml; sp, 25µg/ml; gm, 15µg/ml).

### Method Details

#### Strain and plasmid construction

All oligos, sgRNAs, plasmids, and strains used in this work are described in **Tables S2-5**. WT PCC 7002 is the background genotype for all strains described. To create strains in **Table S5**, noted plasmids, or amplicons containing homologous recombination arms and inserts, were transformed into WT or mutant backgrounds, and transformants were selected on the specified antibiotic(s). To transform, we mixed ∼1 ug of plasmid or amplicon with day old cells and allowed them to incubate for 4-14 hr before plating on 1% Bacto-agar plates with antibiotics. Plates also included 0.5 µg/mL aTC for all *240x tetO-array: TetR-sfGFP* strains. Once individual colonies were detectable, they were patched to new plates with single or combined antibiotics and aTC. After ∼48 hr of growth patches were checked for segregation and intact *tetO* arrays. Strains were considered segregated when no WT products could be detected from PCR using primers flanking the insert and/or gene specific primers (**Table S2**). All CRISPRi strains were freshly transformed for each experiment due to genetic instability of the constructs following repeated passaging. We transformed strains in the following order: 1) 240x tetO array containing construct, 2) TetR-sfGFP containing construct, and 3) mOrange2, sgRNA, or deletion construct. In our system, the strand specificity of the TetR-sfGFP gene had an effect on tetO array stability, with negative strand constructs being more stable than positive stand constructs.

All plasmids described in **Table. S3**, except KAMc0006, were created using gibson assembly of PCR amplified inserts and backbones from base plasmids described in the STAR Methods Resource Table or the PCC 7002 genome. *mOrange2* was subcloned from mOrange2-pBAD, a gift from Michael Davidson, Nathan Shaner, and Roger Tsien (Addgene plasmid #54531). KAMc0006 was created with restriction digest cloning to insert the 240x tetO:GmR array from PRS316-240xtetO into the JCC257 backbone. PRS316-240xtetO was a gift from Narendra Maheshri (Addgene plasmid #44755).

#### Hoechst Staining

Cells were stained with Hoechst dye at specific growth phases. Cells were briefly centrifuged and resuspended in low salt (10% of normal) A+ media and mixed with 5 µg/mL 33342 Hoechst stain (Fisher, CAS # 23491-52-3). Cells were then incubated in the dark for ∼35 min before being washed twice in full salt A+ media. We imaged cells using 395 nm excitation and collected light emitted between 425-477 nm.

#### Quantitative Real Time PCR (qRT-PCR)

To determine average chromosome copy number from bulk culture, we followed the procedure described by Pecoraro et al., (2011). Briefly, we counted either WT or scJC0147 cells at the noted growth phase and then extracted DNA using phenol:chloroform extraction (Green, Sambrook, & Sambrook, 2012) after initial treatment with 5 mg/mL lysozyme for 1hr shaking prior to SDS/proteinase K lysis. To create samples for a standard curve, we purified and quantified ∼1000 bp PCR products amplified from the PCC7002 genome. These fragments were serially diluted to create standards of known concentrations. We used the ThermoFisher QuantStudio6 platform to perform qRT-PCR. Samples were prepared with PerfeCTa SYBR Green SuperMix Reaction Mix (QuantasBio, Catalog # 95056-500) and primers that annealed within our 1000 bp standards (**Table S2**). Samples were run in triplicate to control for pipetting error. We determined average chromosome copy number by measuring the amount of DNA in each sample and dividing by original cell count. Averages and standard deviations were calculated from 2-3 biological replicates of each strain in each condition.

#### sgRNA design and CRISPR-interference

sgRNAs described in **Table S4** were designed using the CRISPy-web platform (Blin, Pedersen, Weber, & Lee, 2016). sgRNAs were chosen based on guidelines described by Gordon et al., (2016). To create an inducible sgRNA construct that was compatible with our chromosome labeling, we placed the sgRNA downstream of the Isopropyl β -D -thiogalactopyranoside (IPTG) inducible cLac94 promoter (Markley et al., 2015). 5 mM IPTG (Fisher, CAS # 367-93-1) was used to induce sgRNA expression. Strains were maintained in the absence of IPTG.

#### Quantitative Long-term Timelapse microscopy

We used a customized Nikon TiE inverted wide-field microscope setup equipped with a Near-IR-based Perfect Focus system, a custom Lexan environmental enclosure for temperature and CO_2_ control, an individually controllable RGB LED light source for transillumination (Lida Light Engine, Lumincor, Beaverton, OR), a high-speed light source with customized filter sets for imaging (Spectra X Light Engine, Lumencor, Beverton, OR), and a synchronized and hardware-triggered motorized shutter for light control for all microscopy described. We specifically used 395 nm 470 nm, 555 nm, and 640 nm excitation wavelengths. Emissions were collected using standard BFP, FITC, mOrange2, or Cy5 filters. Images were acquired on an ORCA Flash4.0 V2+ Digital sCMOS camera (Hamamatsu) and a Nikon CFI60 Plan Apochromat Lamda 100x oil immersion objective (1.45 N.A.). We used NIS Elements Software with Jobs acquisition upgrade for image capture. Fiji (Schindelin et al., 2012) was used to crop and reformat images for publication.

For either single frame or time-lapse microscopy, we spotted ∼2 µL of exponential-phase cells on A+ media pads (1% agarose w/v) and allowed them to air dry before inverting them on to 35 mm glass bottom dishes (Ibidi). Unless otherwise noted, all long-term imaging of strains with the *tetO* array was performed with 0.05 µg/mL aTC added to the imaging pad prior to solidification. Additionally, we added 5mM IPTG to induce sgRNA expression, 1.25 µg/mL rifampicin (Fisher, CAS# 13292-46-1) to inhibit transcription, or 3.4 µg/ml chloramphenicol (Fisher, CAS# 591-50-4) to inhibit translation in noted experiments. For CRISPRi induction as well as rifampicin and chloramphenicol treatment, imaging pads were placed at 30°C for ∼1hr before cells were spotted on to pads. Imaging was started ∼15 min after spots air dried. For all other imaging, cells were acclimated to imaging conditions (30°C and noted 640 nm transmitted light intensity) for ∼1 hr prior to initial imaging. Images were taken at a 30 min frame rate for all experiments, except those described in **Fig. 6F-I**, where the frame rate was 10 min. Cells were continuously illuminated with 640nm transmitted light except during fluorescent imaging.

#### Image Processing and Analysis

Image processing was carried out using custom MATLAB programs. From initial testing, we found that segmentation was most reliable on brightfield images of the cell, but with the focus offset by 2 µm. In brief, cell identification (segmentation) was performed by thresholding this brightfield offset image. The threshold level was chosen by computing the intensity histogram of the image. Since the images contained both dark background and bright cells, the intensity histogram appeared bimodal, allowing the peak of the background intensity distribution to be identified. A Gaussian model was then fit to this background intensity. The threshold intensity was then chosen to be the mean + *F* * standard deviation of the fitted Gaussian. The threshold factor, *F*, was optimized for each set of images, and ranged from 2.0-3.5.

Once the threshold intensity was determined, an initial binary mask was created by setting pixels in the image which were brighter than the threshold to true, and all other pixels to false. This initial mask often contained connected cells that were physically close to each other. Individual cells were then separated using the watershed algorithm. Data from individual cells were then tracked over time using the linear assignment algorithm (Jaqaman et al., 2008). When necessary cell masks were corrected by hand to avoid obvious tracking and/or size errors.

To count chromosomes in the images, sfGFP labeled puncta were identified using the difference of Gaussians method (Lowe, 1999). Here, two Gaussian blur filters were applied to the original image, with standard deviations σ of 1.12 and 1.58 pixels. The difference between the two images was then computed, and puncta were identified by intensity thresholding. Individual puncta were then counted to obtain chromosome numbers in each cell.

### Quantification and Statistical Analysis

During growth, the length of cells was assumed to be increasing exponentially. Hence, we obtained a growth rate by fitting the log of cell length over time to the linear function

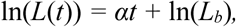

where *L*(*t*) is the length of the cell at time *t, α* is the growth rate, and *L*_*b*_ is the length at birth. Doubling times were calculated by dividing ln(2) by the growth rate *α.*

#### Corrected Chromosome Counting Algorithm

Because chromosome labeling is a dynamic process, we developed an algorithm based on both previous and future chromosome counts to create a more accurate representation of chromosome number during time lapse imaging. The following rules were used to define chromosome number:

1. Chromosome number could not decrease
2. Chromosome number only increased in the *nth* frame if the chromosome number in the *n+1* frame was equal to or greater than the number recorded in the *nth* frame
3. For the final frame of a cell trace chromosome number only increased if the sum of chromosome counts in the two daughter cells was equal to or greater than the number recorded in the final frame

Representations of original and corrected chromosome counts are displayed in **Fig S2**.

#### Data and Statistics

The length of each movie, as well as number of cells analyzed in each experiment, is recorded in the figure legends. For cells with measured growth rates, only those with < 0.3 norm of residuals between the data and the model were kept in the analysis. All long-term imaging experiments were performed at least twice to ensure reproducible phenotypes. However, only data from a single long-term imaging experiment is shown with the exception of Fig. 5G-I to ensure that enough cells were analyzed.

Chromosome counts were verified by eye for 50-100 cells for each still or time lapse imaging experiment. For time-lapse experiments, chromosome counts were checked at multiple stages of imaging. Images displayed in figures have been smoothed to reduce background noise. However, all analysis was done on the original images. To normalize for variations in imaging cell intensity, measurements were smoothed over 120min segments in **Fig 6G** (bottom panel) and **I**.

For **Fig. 3I-K** and **P-Q** cells were categorized as growing predominantly in HL if they grew for at least 3 hr in HL conditions. Cells were categorized as growing predominantly in LL if they grew for at least 7 hr in LL conditions. To avoid grouping cells that grew in both light conditions cells in either group were analyzed only if they grew for less than 1hr in the alternative light condition.

All boxplots denote the median (light gray line), the 25^th^ and 75^th^ quartiles of the data (dark gray boxes), and the most extreme values not considered outliers (whiskers). Whisker values correspond to approximately +/-2.7 σ and represent ∼99.3% of the data, assuming normal distribution. For all histogram and single frame data, chromosome counts are only displayed for cells within 3 standard deviations of the mean of the data. To determine if the variation between cell populations was statistically significant, we used the two sample Kolmogorov-Smirnov test. Significance was determined using Bonferroni corrected *p*-values, which are denoted by *, * *, * ** to represent <0.01, 0.001, and 0.0001, respectively.

### Data and Software Availability

The code developed to segment and quantify images was custom generated for this work and is available at: https://biof-git.colorado.edu/cameron-lab-public/multigenanalysis-polyploid-moore

## SUPPLEMENTAL INFORMATION

**Fig S1.**
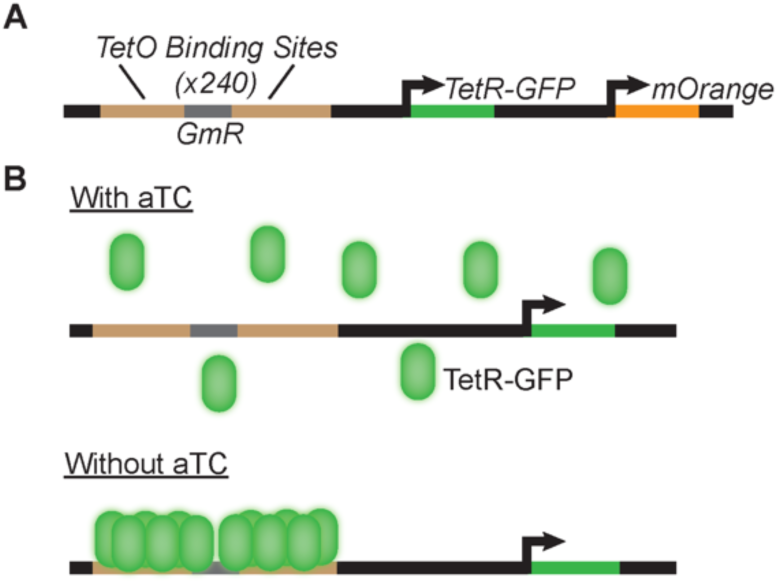
Schematic of tetO array:TetR-sfGFP chromosome labeling system (related to Fig 1). **A)** Depiction of *mOrange2::240x tetO array::TetR-sfGFP* genotype. sfGFP = super folder GFP. **B)** Schematic of TetR-sfGFP binding to the tetO array with (top) and without (bottom) aTC.

**Fig S2.**
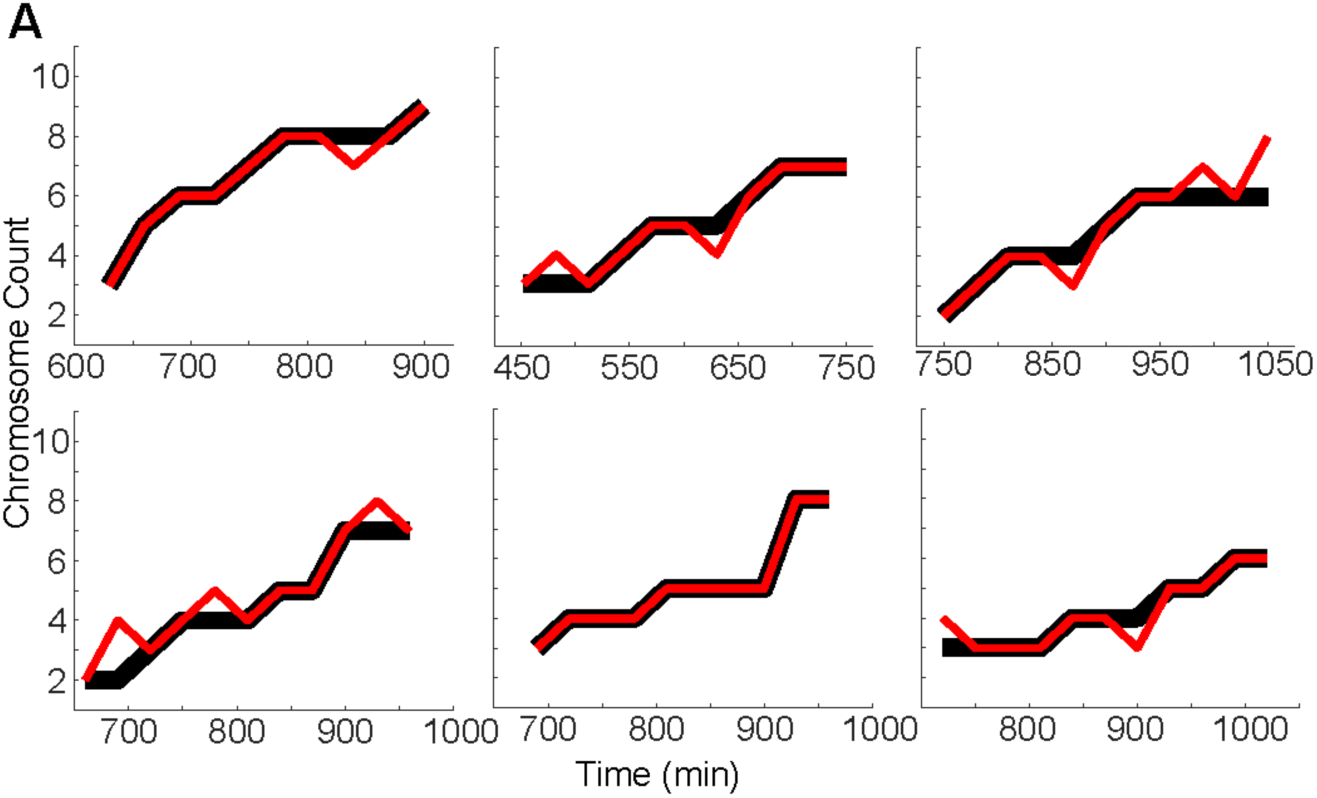
Representative traces of corrected and raw chromosome counts over time (related to Fig 2). A) Raw chromosome counts are traced in red, corrected chromosome counts are traced in black. The X-axis represents time-elapsed from the first frame of time lapse imaging. See Methods Details for information regarding the chromosome correcting algorithm used.

**Fig S3.**
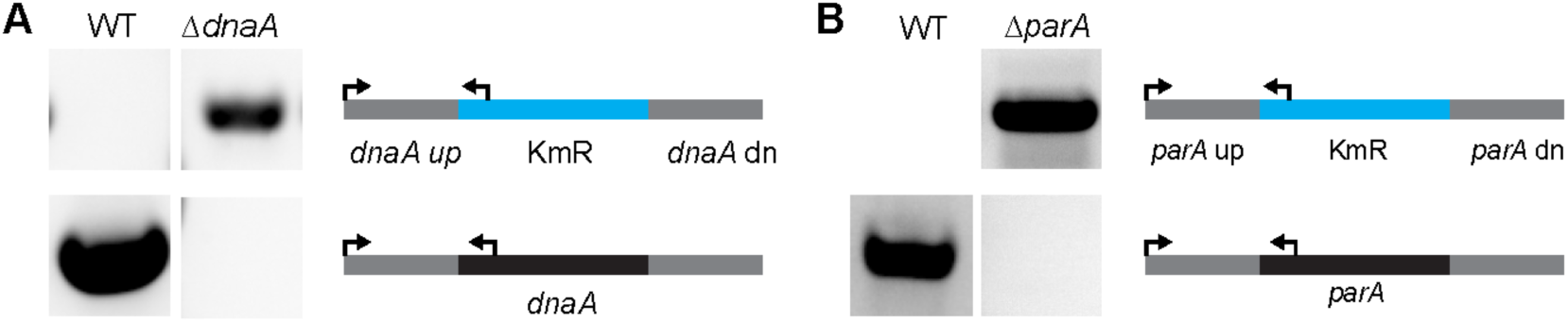
Analysis of of *ΔdnaA* and *ΔparA* strains (related to Figs. 5 and 6). **A)** PCR products confirming presence of *KmR* insert at the *dnaA* locus (top panel) and absence of WT product (bottom panel). **B)** PCR products confirming presence of *KmR* insert at the *parA* locus (top panel) and absence of WT product (bottom panel).Schematics not to scale.

## Supplemental Tables

**Table S1.** Average chromosome counts from bulk culture (related to Fig 1). Average chromosome counts from bulk culture *mOrange2::240x tetO array::TetR-sfGFP* (scJC0147) or WT PCC 7002 cells grown to exponential or stationary phase. *mOrange2::240x tetO array::TetR-sfGFP* cultures were resupplied with 0.5µg/mL every 48hr.

| Strain | Growth Phase | Average Chromosome Number | Standard Deviation |
| --- | --- | --- | --- |
| scJC0147 | Exponential | 3.8 | +/- 0.4 |
| scJC0147 | Stationary | 1.8 | +/- 0.3 |
| PCC 7002 | Exponential | 3.3 | +/- 0.6 |
| PCC 7002 | Stationary | 2.3 | +/- 0.1 |

**Table S2.**
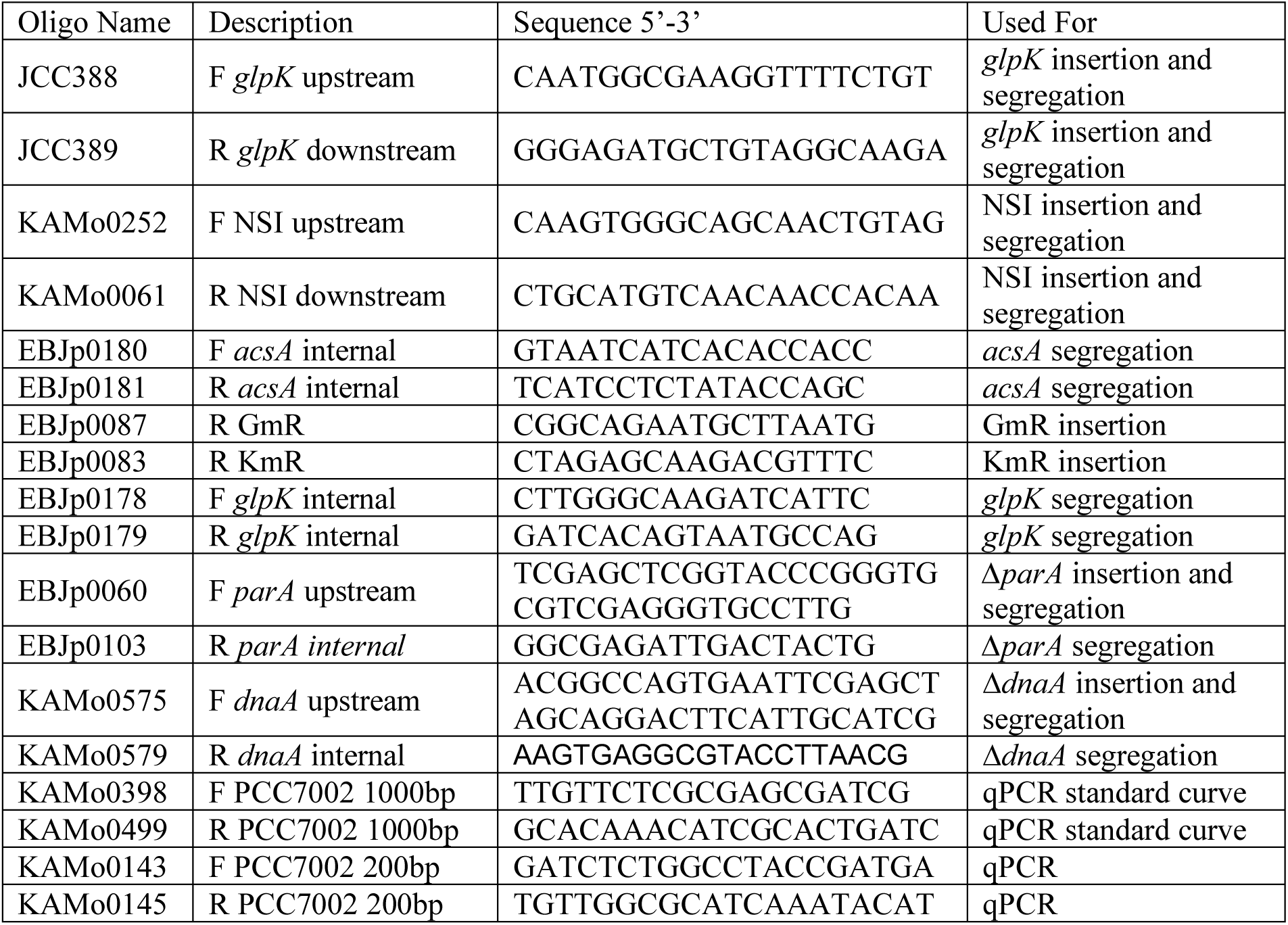
Strain Segmentation and qPCR Oligonucleotides. PCC 7002 gene names are italicized. NSI is a neutral site described by Davies, Work, Beliaev, & Posewitz, 2014)

**Table S3.**
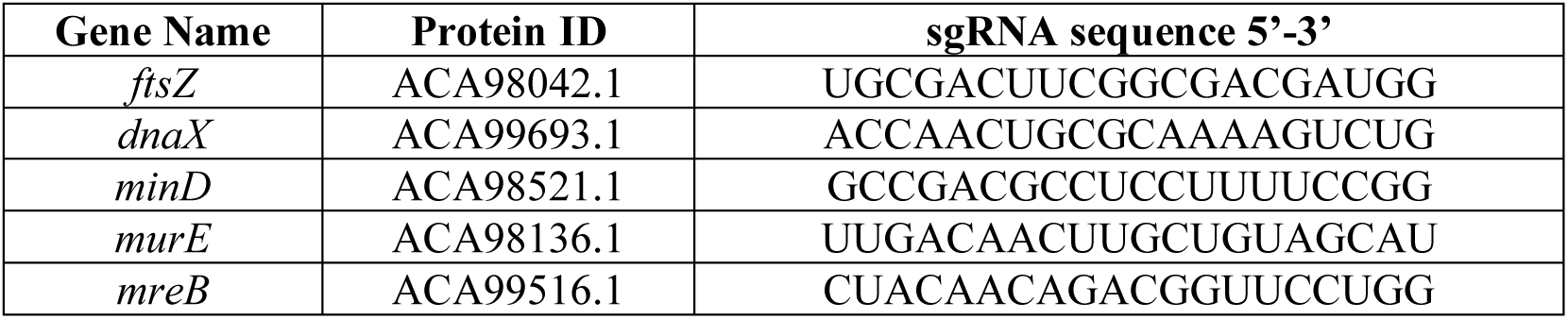
Gene name, protein ID, and sgRNA sequences.

**Table S4.** Plasmids used in this study. GmR = gentamycin resistance. KmR = kanamycin resistance, SpR = spectinomycin resistance. NSI is a neutral site described by Davies et al., 2014. PCC7002 gene names are italicized.

| Plasmid Name | Description | Neutral Site | Reference |
| --- | --- | --- | --- |
| KAMc0006 | 120x tetO-array flanking GmR Cassette | *glpK* | This work |
| KAMc0064 | pJ23113:tetR-sfGFP:SpR cassette | *acsA* | This work |
| KAMc0084 | pJ23113:tetR-sfGFP:dCas9:SpR cassette | *acsA* | This work |
| KAMc0131 | pCpt:mOrange2:KmR cassette | NSI | This work |
| KAMc0189 | pJ23119:mOrange2:KmR cassette | NSI | This work |
| KAMc0197 | *dnaA* upstream and downstream regions flanking KmR cassette | N/A | This work |
| KAMc0205 | cLac94:ftsZ-sgRNA3:LacI:KmR cassette | NSI | This work |
| KAMc0208 | cLac94:dnaX-sgRNA:LacI:KmR cassette | NSI | This work |
| KAMc0211 | cLac94:minD-sgRNA:LacI:KmR cassette | NSI | This work |
| KAMc0217 | cLac94:murE-sgRNA:LacI:KmR cassette | NSI | This work |
| KAMc0223 | cLac94:mreB-sgRNA2:LacI:KmR cassette | NSI | This work |
| EBJc0022 | *parA* upstream and downstream regions flanking KmR cassette | N/A | This work |

**Table S5.** PCC7002 derived strains. Plasmid descriptions can be found in Table S4. GmR - gentamycin resistance, Gm - gentamycin, KmR - kanamycin resistance, Km - kanamycin, SpR - spectinomycin resistance, Sp - spectinomycin. Genotypes are italicized.

| Strain Name | Description | Transformed Plasmids | Ab Resistance |
| --- | --- | --- | --- |
| Wild Type (WT) | *Synechococcus* PCC 7002 | None | None |
| scJC0134 | $\Delta glpK:240x\ tetO\ array:GmR :: \Delta acsA:tetR-sfGFP:SpR$ | KAMc0006, KAMc0064 | Gm, Sp |
| scJC0147 | $\Delta glpK:240x\ tetO\ array:GmR :: \Delta acsA:tetR-sfGFP:SpR::\Delta NSI:pCpt-mOrange2:KmR$ | KAMc0006, KAMc0064, KAMc0131 | Gm, Sp, Km |
| scJC0164 | $\Delta glpK:240x\ tetO\ array:GmR :: \Delta acsA:tetR-sfGFP:dCas9:SpR$ | KAMc0006, KAMc0084 | Gm, Sp |
| scJC0182 | $\Delta glpK:240x\ tetO\ array:GmR :: \Delta acsA:tetR-sfGFP:SpR::\Delta NSI:pJ23119-mOrange2:KmR$ | KAMc0006, KAMc0064, KAMc0189 | Gm, Sp, Km |
| scJC0189 | $\Delta glpK:240x\ tetO\ array:GmR :: \Delta acsA:tetR-sfGFP:SpR::\Delta dnaA:KmR$ | KAMc0006, KAMc0064, KAMc0197 | Gm, Sp, Km |
| KAMs0003 | $\Delta glpK:240x\ tetO\ array:GmR :: \Delta acsA:tetR-sfGFP:SpR::\Delta parA:KmR$ | KAMc0006, KAMc0064, EBJc0022 | Gm, Sp, Km |

## Supplemental Movie Legends

**Movie S1. Time lapse imaging of labeled chromosomes in rapidly growing cells (related to Fig 2).** *mOrange2::240x tetO array::TetR-sfGFP* cells were grown with 150 µmol photons m^−2^ s^−1^ red light on 1% agarose pads supplemented with 0.05 µg/mL aTC for 18.5 hr. sfGFP labeled chromosomes shown. For all supplemental movies scale bar = 5 µm.

**Movie S2. Time lapse imaging of ftsZ CRISPRi cells (related to Fig. 4)**. *ftsZ sgRNA::240x tetO array::TetR-sfGFP* cells were grown with 150 µmol photons m^−2^ s^−1^ red light on 1% agarose pads supplemented with 0.05 µg/mL aTC and 5 mM IPTG for 21.5 hr. Left panel: sfGFP labeled chromosomes, Right panel: brightfield imaging (imaged with 640 nm transmitted light). White dot in lower left corner indicates frames that were *not* analyzed due to difficulty segmenting cells and identifying chromosomes due to overlapping cell growth.

**Movie S3. Time lapse imaging of murE CRISPRi cells (related to Fig. 4)**. *murE sgRNA::240x tetO array::TetR-sfGFP* cells were grown with 150 µmol photons m^−2^ s^−1^ red light on 1% agarose pads supplemented with 0.05 µg/mL aTC and 5 mM IPTG for 21.5 hr. Left panel: sfGFP labeled chromosomes, Right panel: brightfield imaging (imaged with 640nm transmitted light).

**Movie S4. Time lapse imaging of dnaX CRISPRi cells (related to Fig. 5).** *dnaX sgRNA::240x tetO array::TetR-sfGFP* cells were grown with 150 µmol photons m^−2^ s^−1^ red light on 1% agarose pads supplemented with 0.05 µg/mL aTC and 5 mM IPTG for 26 hr. sfGFP labeled chromosomes shown.

**Movie S5. Time lapse imaging of mreB CRISPRi cells (related to Fig. 6).** *mreB sgRNA::240x tetO array::TetR-sfGFP* cells were grown with 150 µmol photons m^−2^ s^−1^ red light on 1% agarose pads supplemented with 0.05 µg/mL aTC and 5 mM IPTG for 15.5 hr. sfGFP labeled chromosomes shown.

**Movie S6. Time lapse imaging of minD CRISPRi cells (related to Fig. 6).** *minD sgRNA::240x tetO array::TetR-sfGFP* cells were grown with 150 µmol photons m^−2^ s^−1^ red light on 1% agarose pads supplemented with 0.05 µg/mL aTC and 5 mM IPTG for 19.5 hr. sfGFP labeled chromosomes shown.

**Movie S7. Time lapse imaging of stationary phase cells with aTC (related to Fig. 6).** *mOrange2::240x tetO array::TetR-sfGFP* cells grown to stationary phase in shaking flasks were grown with 150 µmol photons m^−2^ s^−1^ red light on 1% agarose pads supplemented with 0.05 µg/mL aTC for 24 hr. sfGFP labeled chromosomes shown.

**Movie S8. Time lapse imaging of stationary phase cells without aTC (related to Fig 6).** *mOrange2::240x tetO array::TetR-sfGFP* cells grown to stationary phase in shaking flasks were grown with 150 µmol photons m^−2^ s^−1^ red light on 1% agarose pads without aTC for 65 hr. Every frame represents 2 hr of imaging. Left panel: sfGFP labeled chromosomes, Right panel: endogenous fluorescence emitted after 640 nm excitation. Calibration bar represents intensities between 75-2450 A.U.

